# Identification of Pre-Existing Adaptive Immunity to Cas9 Proteins in Humans

**DOI:** 10.1101/243345

**Authors:** Carsten T. Charlesworth, Priyanka S. Deshpande, Daniel P. Dever, Beruh Dejene, Natalia Gomez-Ospina, Sruthi Mantri, Mara Pavel-Dinu, Joab Camarena, Kenneth I. Weinberg, Matthew H. Porteus

## Abstract

The CRISPR-Cas9 system has proven to be a powerful tool for genome editing, allowing for the precise modification of specific DNA sequences within a cell. Many efforts are currently underway to use the CRISPR-Cas9 system for the therapeutic correction of human genetic diseases. The most widely used homologs of the Cas9 protein are derived from the bacteria *Staphylococcus aureus* (*S. aureus*) and *Streptococcus pyogenes* (*S. pyogenes*). Based on the fact that these two bacterial species cause infections in the human population at high frequencies, we looked for the presence of pre-existing adaptive immune responses to their respective Cas9 homologs, SaCas9 (*S. aureus* homolog of Cas9) and SpCas9 (*S. pyogenes* homolog of Cas9). To determine the presence of anti-Cas9 antibodies, we probed for the two homologs using human serum and were able to detect antibodies against both, with 79% of donors staining against SaCas9 and 65% of donors staining against SpCas9. Upon investigating the presence of antigen-specific T-cells against the two homologs in human peripheral blood, we found anti-SaCas9 T-cells in 46% of donors. Upon isolating, expanding, and conducting antigen re-stimulation experiments on several of these donors’ anti-SaCas9 T-cells, we observed an SaCas9-specific response confirming that these T-cells were antigen-specific. We were unable to detect antigen-specific T-cells against SpCas9, although the sensitivity of the assay precludes us from concluding that such T-cells do not exist. Together, this data demonstrates that there are pre-existing humoral and cell-mediated adaptive immune responses to Cas9 in humans, a factor which must be taken into account as the CRISPR-Cas9 system moves forward into clinical trials.

## Introduction

Genome editing, a method used to precisely alter the DNA sequence of a cell, is broadly used as a research tool and is now being developed to create gene therapies to treat human disease. The Cas9/guide RNA (gRNA) platform, derived from the CRISPR/Cas9 adaptive bacterial immune system, has proven to be a powerful tool for genome editing because it is relatively simple to use, has high on-target activity, and has high specificity (*1, 2*). Genome editing is initiated by the induction of a site-specific DNA double-stranded break. The Cas9/gRNA system consists of the site-specific Cas9 nuclease that, when complexed with a short gRNA, can be directed to create a double-strand break in a wide variety of DNA sequences in both eukaryotic and prokaryotic genomes (*1, 3*). The ability of the Cas9/gRNA system to create site-specific double-strand breaks in DNA can be applied to treat disease through the knockout of target genes (by non-homologous end-joining (NHEJ)) or homologous recombination (HR)), the deletion of target stretches of DNA (by NHEJ), or the incorporation of new desired DNA sequences into the genome (by HR). These techniques are being applied to create novel cell therapies, either through *ex vivo* editing of cells (*4–6*) followed by the transplantation of engineered cells into a patient, or through *in vivo* editing of the patient’s cells by delivery of Cas9/gRNA via viral vectors (mainly with recombinant adeno-associated vectors (rAAV)) or nanoparticles (*7–9*). For instance, significant progress has been made in using the Cas9/gRNA system to edit hematopoietic stem cells *ex vivo* to treat sickle cell disease (*4, 6*). Similarly, the Cas9/gRNA system has been used to edit cells *in vivo* to address conditions such as muscular dystrophy and retinitis pigmentosa (*10–13*).

Although a variety of Cas9 homologs have been characterized, the most well-developed systems are derived from *Staphylococcus aureus* (*S. aureus*) and *Streptococcus pyogenes* (*S. pyogenes*). The S. *aureus* Cas9 (SaCas9) is primarily applied for *in vivo* editing of cells because it can be more readily packaged into AAV vectors (*8*). The S. *pyogenes* Cas9 (SpCas9) has been shown to have great therapeutic potential in proof-of-concept pre-clinical studies both *ex vivo* and *in vivo* (*4–6, 8, 9*).

*S. aureus* and *S. pyogenes* are common human commensals that can also be pathogenic. 40% of the human population is colonized by S. *aureus* (*14*) and 20% of school-aged children are colonized with S. *pyogenes* at any given time (*15*). Furthermore, studies of humoral responses to these bacteria have found that antibodies against S. *aureus* and S. *pyogenes* are present in 100% of human adults (*16–18*). Similarly, T-cells specific against antigens from these bacteria can be found at a high frequency within the human population (*18, 19*). The abundance of S. *aureus* and S. *pyogenes* within the human population, as well as widespread adaptive humoral and cell-mediated immune responses to both species, raises the possibility that humans may also have pre-existing adaptive immunity to the most commonly used homologs of Cas9 that are derived from these bacteria. Prior work, in which recombinant S. *aureus* Cas9 was expressed in mice, found a clear adaptive immune response to Cas9, indicating that mammalian immune systems recognize Cas9 as an antigen (*20*).

The presence of pre-existing adaptive immune responses in humans to either Cas9 homolog may hinder the safe and efficacious use of the Cas9/gRNA system to treat disease, and may even result in significant toxicity to patients. In prior gene therapy trials in which patients had pre-existing adaptive immune responses to the viral vector, no therapeutic benefit was derived from the therapy due to either neutralization of the vector by antibodies (*21, 22*) or clearance of transduced cells by cytotoxic T-lymphocytes (CTLs) (*23*). Similarly, there have been rare cases in gene therapy trials in which a patient has pre-existing adaptive immunity to the delivered transgene, resulting in a lack of any therapeutic effect from treatment and an expansion of transgene-product-specific T-cells (*24*). Adaptive immune responses can also induce systemic inflammatory responses, resulting in serious toxicity. The gene therapy field suffered a significant setback when a single patient died from a systemic inflammatory response to a gene therapy vector(*25*). Thus, the human adaptive immune response can be a tremendous barrier to both safety and efficacy of *in vitro* and *in* vivo gene therapy and cannot be reliably evaluated using non-human systems(*26*). It follows, then, that a pre-existing adaptive immune response to Cas9, whether humoral or cellular, raises similar issues as the Cas9/gRNA system enters use in clinical trials. For example, a CTL response against Cas9 would result in the destruction of any cells that are presenting Cas9 peptides on their MHC molecules, potentially eliminating edited cells and rendering the therapy ineffective(*27*).

Based on the widespread presence of adaptive immunity to the bacteria from which SaCas9 and SpCas9 are derived, we hypothesized that there might be pre-existing humoral and cell-mediated responses to Cas9 in humans. Our results demonstrate that there is pre-existing humoral immunity to both SaCas9 and SpCas9; that there is clear evidence of a cellular immune response to SaCas9 in some healthy individuals; and that although we did not detect it, we cannot rule out that there may also be pre-existing cellular immunity to SpCas9.

## Results

### Detection of IgG to Cas9 from Healthy Humans

We first investigated if there was a pre-existing humoral response to Cas9. To search for pre-existing IgG antibodies against SaCas9 and SpCas9, we used serum from human cord blood to probe for Cas9 by immunoblot. Because IgG crosses the placental barrier, the IgG content of cord blood reflects the seroprevalence of an IgG antibody in the mother. We found that IgG antibodies to both SaCas9 and SpCas9 could be detected in human serum (Fig. 1). The specificity of the antibody against Cas9 was demonstrated by showing immunoreactivity to CD34+ cells that had been electroporated with SaCas9 and SpCas9, but not to CD34+ cells that had not been electroporated (Supplementary figure 1). Using Western blotting to detect seroreactivity against purified Cas9 among 22 cord blood donors, we found that at a serum dilution of 1:10, 86% of donors stained positively for SaCas9 and 73% for SpCas9, with some donors staining for one homolog of Cas9 but not the other (Table 1). Having found that we could detect antibodies against Cas9 in cord blood donors at a high frequency, we then probed for Cas9 with serum from peripheral blood drawn from 12 healthy adults. Probing for SaCas9 and SpCas9 with serum at a dilution of 1:10, we detected anti-SaCas9 antibodies among 67% of donors and anti-SpCas9 among 42% (Table 1). The discrepancy between the frequencies of seroprevalence in the cord blood as compared from peripheral blood could be the result of multiple factors, including that the two sources of serum were processed in different ways and that the IgG antibody might be sensitive to the method of processing.

**Figure 1:**
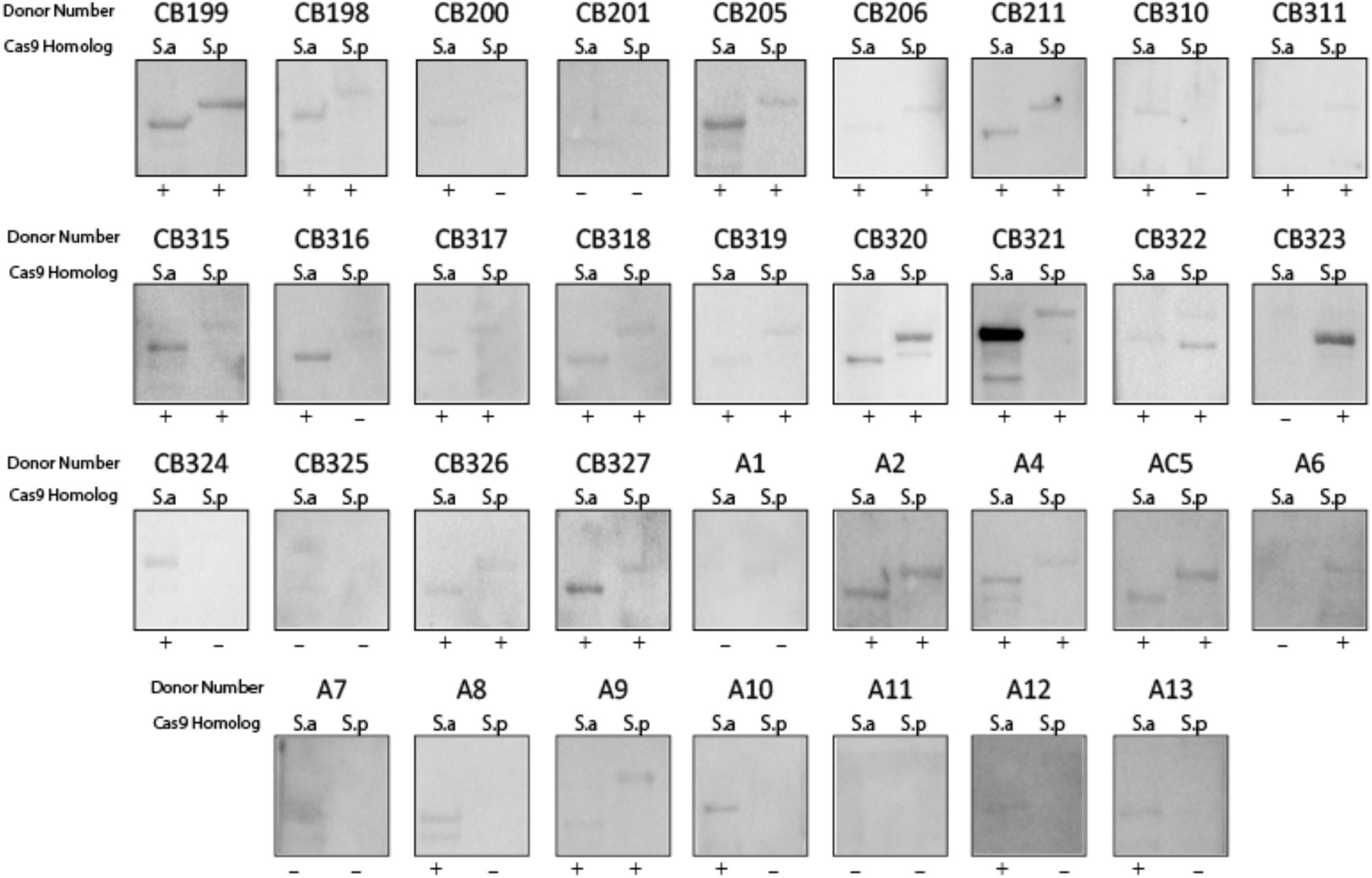
Identification of Pre-Existing Humoral Immunity to Cas9 Protein. Western blot analysis using healthy donor serum to determine if humans have IgG antibody to Cas9 proteins. Shown are serum either derived from cord blood (labelled “CB”) or adult (labelled “A”) and the immunoreactivity at 1:10 dilution against purified Cas9 proteins (S.a for S. aureus Cas9 and S.p for S. pyogenes Cas9). Each blot was scored as either positive (+) or negative (−) for immunoreactivity.

**Table 1:** Frequencies of Humoral and Cellular Immunoreactivity to Cas9 Protein

| Cas9 Homolog | Number of Positive Cord Sera | Number of Positive Adult Sera | Number of Negative Cord Sera | Number of Negative Adult Sera | Seropositive | Number of Healthy Donor PBMCs Analyzed | Number (percent) Healthy Donor PBMCs positive by IFN-γ Capture |
| --- | --- | --- | --- | --- | --- | --- | --- |
| Aureus | 19 | 8 | 3 | 4 | 79% | 13 | 6 (46%) |
| Pyogenes | 16 | 5 | 6 | 7 | 65% | 13 | 0 (0%) |

The presence of anti-Cas9 antibodies demonstrates that the human immune system is exposed to and generates specific humoral responses against SaCas9 and SpCas9. The presence of anti-Cas9 antibodies reveals that underlying mechanisms exist in the human immune system for B-cell recognition of and antibody production after exposure to Cas9, likely through interaction with T_H_2 CD4+ T-cells (*28*).

### Detection of Antigen Specific T-cells to SaCas9

The finding of humoral anti-Cas9 responses prompted us to explore if Cas9-specific T_H_1 CD4+ T-cells and corresponding CTLs also existed in humans.

To determine if there were antigen-specific T-cells against SaCas9 and SpCas9, we used a cytokine capture system. In the cytokine capture system (CCS), an antigen is added to a mixture of peripheral blood mononuclear cells (PBMCs). This antigen is spontaneously taken up by antigen presenting cells (APCs), processed, and then expressed as MHC-peptide complexes on the APC surface. Presentation of the peptides to antigen-specific T-cells activates them, prompting them to secrete the inflammatory cytokine interferon-γ (IFN-γ) (*29*). IFN-γ is transiently present on the cell surface of T-cells as secretory vesicles fuse with the plasma membrane. Antigen-specific T cells expressing IFN-γ can be detected, phenotyped, and then isolated by anti-IFN-γ antibodies (Fig. 2A).

**Figure 2:**
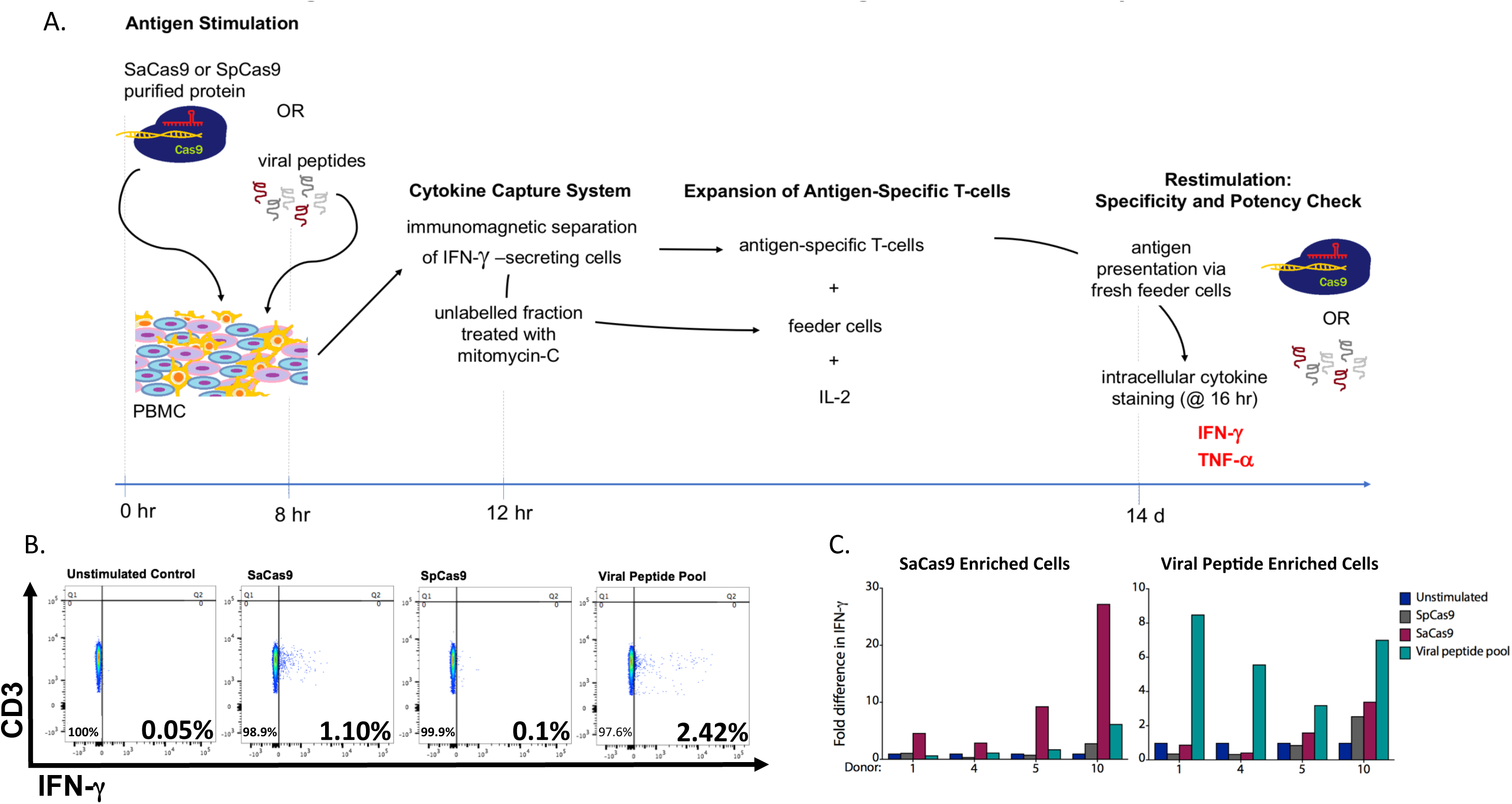
Identification of Pre-Existing T-cell Reactivity to Cas9. A) Workflow for the identification, maintenance, and authentication of T-cells directed against Cas9 homologs or viral peptides. B) Representative flow cytometry plots of IFN-γ production by live T-cells during initial screen of healthy donor PBMCs using cytokine capture system (see Supplementary Fig. 2 for gating strategy). This donor was marked as responsive to SaCas9 but not SpCas9 and was responsive to the positive control viral peptide pool. C) Plots showing fold difference in intracellular IFN-γ production by SaCas9-specific (left) and viral peptide-specific (right) T-cells for each donor, as compared to donor’s unstimulated baseline, in various antigen re-challenge conditions.

PBMCs from 13 healthy adult donors were incubated with SaCas9 protein, SpCas9 protein, or a mixture of 17 antigenic viral peptides, which served as a positive control. Cells were then isolated by CCS and characterized by flow cytometry for surface marker expression to identify IFN-γ secreting T-cells (Supplementary figure 2A). A donor was considered positive for having a T-cell response against a Cas9 homolog if their PBMCs had at least a five-fold increase in the number of T-cells secreting IFN-γ as compared to T-cells from an unstimulated sample of their PBMCs (Fig. 2B). We detected IFN-γ responses from both CD4+ and CD8+ T-cells directed against SaCas9 in 46% of donors (Table 1, Supplementary figure 2B). In some experiments, some donor PBMCs showed IFN-γ response to SpCas9, but this response was not consistent between replicates and we have conservatively scored all samples as negative for having antigen-specific T-cells directed against SpCas9 (data not shown). In addition, stimulated T-cells were found to have higher expression of other activation markers, such as CD69, CD154, and CD25 (Supplementary figure 3A) (*30–32*), as compared to the unstimulated baseline. As a negative control, we also stimulated human cord blood T-cells, which have no previous exposure to antigens, with the two Cas9 homologs and the viral peptide library (n = 2). We found that T-cells in these samples did not have an IFN-γ response to any antigen, as expected based on the naïve cellular immune system of a newborn (Supplementary figure 4), confirming that the T-cell response in healthy adult donor samples was a result of immune responses that occurred due to postnatal exposure and not due to activation upon first *in vitro* exposure to the antigen.

We analyzed whether donors who had T-cell responses also had IgG antibody against Cas9. We found that 71% of donors who had a T-cell response against SaCas9 also had antibodies against SaCas9, as detected by immunoblot. This suggests that many donors who have antigen-specific T-cells that recognize SaCas9 also have T_H_2 CD4+ cells and B-cells that recognize SaCas9.

As a control to determine that the T-cell response we were detecting against SaCas9 was antigen-specific, we selected four PBMC donors with an IFN-γ response against SaCas9 for antigen-specific T-cell enrichment, expansion, and re-stimulation (Fig. 2A). IFN-γ-secreting T-cells were magnetically separated from the stimulated PBMC population and cultured on autologous feeder cells. The cultured cells were mostly T-cells: few to no B-cells or monocytes present, and both CD4+ and CD8+ T-cells were isolated and were present in the cultured population (Supplementary figure 5). The ratio of these two T-cell subpopulations for a given donor sample was found to be dependent on the IL-2 concentration in the media, so the IL-2 concentration was chosen to maintain a distribution of both CD4+ and CD8+ T cells.

To determine if the enriched and expanded T-cells were antigen-specific, we checked their specificity and potency via intracellular cytokine staining following re-exposure to antigen after approximately two weeks of culture (Fig. 2A). The experiments included a cross-stimulation, in which antigen-specific T-cells were re-challenged not only with their respective antigens, but also with other antigens to identify cross-reactive or non-specific responses.

In 4 out of 4 donors from whom SaCas9-specific T-cells were expanded, activated (CD69+) SaCas9-specific T-cells showed a higher fold-increase in intracellular IFN-γ production when re-stimulated with SaCas9, as compared to the unstimulated baseline, and as compared to re-stimulation with SpCas9 or the viral peptide pool (Fig. 2C, Supplementary figure 6A). 3 out of 4 donors showed a similar pattern of intracellular TNF-α production upon re-stimulation (Supplementary figure 7). Similarly, our positive control, the antigen-specific T-cells directed against the viral peptide pool, responded exclusively to the viral peptide pool upon re-stimulation, confirming the specificity of our assay (Fig 2C, Supplementary figure 6B).

Upon intracellular cytokine staining, re-challenged antigen-specific T-cells were found to have higher expression of activation markers CD69, CD154, and CD25 (Supplementary Figure 3B). In our analysis, we focused on CD69+ T-cells, as CD69 is an early T-cell activation marker(*31*). We also considered primarily anti-SaCas9 T-cells’ ability to produce IFN-γ upon re-stimulation, as T-cell clones isolated via the cytokine capture system were initially selected based on their IFN-γ production ability. It is possible that the isolated cells vary in their ability to also produce TNF-α (*33*), so not all donors’ antigen-specific T-cell responses included significantly higher TNF-α production above other experimental conditions.

Together, this data demonstrates that many healthy humans have pre-existing antigen-specific T-cells directed against SaCas9, and that these cells can undergo expansion and can express effector molecules. We were unable to detect, expand, and re-stimulate antigen-specific T-cells directed against SpCas9 using this cytokine capture assay.

## Discussion

In this study, we provide evidence of pre-existing adaptive immune responses to Cas9 in humans, which probably develop after postnatal exposure. We detected IgG antibodies against both SaCas9 and SpCas9 in the population and were also able to detect antigen-specific T-cells directed against SaCas9. This data raises a potential barrier to the safe and efficacious use of the Cas9/gRNA system to treat disease.

Although we detected IgG antibodies to both Cas9 homologs in human serum via immunoblotting, the finding of IgG antibodies against Cas9 may not be detrimental to the clinical use of Cas9. Currently, the primary methods to deliver Cas9 into cells involve its direct transfer into cells using either *ex vivo* methods, such as electroporation(*4–6*), or *in vivo* methods, such as viral vectors or nanoparticles(*7–9*). In both of these settings, the Cas9 protein would not be directly exposed to IgG antibodies. As such, antibodies present in human serum against Cas9 may not directly inhibit the efficacy of a Cas9-based therapy. If the Cas9 protein is directly delivered *in vivo*, such as through a ribonucleoprotein (RNP) complex, pre-existing antibodies could potentially decrease effectiveness unless the RNP was protected from IgG binding. Moreover, the apparent widespread presence of antibodies against both SaCas9 and SpCas9 is indicative of the frequency with which adaptive immune responses to Cas9 develop within the human population. In clinical gene therapy trials, patients are excluded from participation if they are found to have pre-existing antibodies to either the vector or the transgene; there is no clinical data about how to safely and effectively administer a gene therapy product to a patient who has pre-existing humoral immunity to one of the components, such as Cas9.

A more significant concern in using Cas9 for clinical applications is the presence of antigen-specific T-cells directed against Cas9. Our data indicates that antigen-specific T-cells, both T_H_1 CD4+ T-cells and cytotoxic CD8+ T-cells, specifically directed against SaCas9, exist within the human population. The SaCas9 T-cells are potent, as indicated by their ability to produce the inflammatory cytokines IFN-γ and TNF-α, both of which are known to play a key function in CTL-mediated clearance of infected cells(*34*). Furthermore, these cells selectively produce high levels of cytokines upon re-exposure to SaCas9, demonstrating that they are antigen-specific. The existence of the pre-existing cellular immunity would likely both limit the effectiveness of and create safety concerns for Cas9-based genome editing. In past gene therapy trials, antigen-specific T-cells against gene therapy vectors and delivered transgenes have eliminated cells that received the therapy, nullifying any therapeutic effect(*23, 24*). It follows, then, that anti-Cas9 CTLs could kill any cells presenting Cas9 peptides on their MHC class I molecules, eliminating edited cells and rendering a therapy ineffective. In addition, anti-Cas9 CTLs could potentially create serious safety problems by inducing a systemic inflammatory response or by eliminating transduced- or nanoparticle-infected cells throughout the body. For example, while AAV2 and AAV8 have been used pre-clinically in mouse models to target the muscle, they also transduce other tissues, such as the liver, and thus one might observe serious hepatic inflammation and hepatocyte death.

Although we were unable to definitively detect antigen-specific T-cells against SpCas9 in this study, this does not necessarily establish that they are not present within the human population. The CCS for detection of antigen-specific T-cells depends on high surface level expression of IFN-γ and may underestimate the frequency of antigen-specific T-cells, as compared to other assays such as ELISpot. Other studies that have attempted to identify antigen-specific T-cells by IFN-γ response have noted that a high frequency of donors, who were eventually determined to have antigen-specific T-cells against an antigen, have gone undetected in an initial screen due to antigen-specific T-cells existing at too low of a frequency to easily detect. This is despite antibodies against the same antigen being detected in the same patient (*23*). The presence of humoral responses against SpCas9 indicates that the healthy human immune system is exposed to and can respond to SpCas9 protein. Further work is required to rule out the existence of SpCas9-specific T-cells.

The presence of antigen-specific T-cells against Cas9 may prove to be most problematic for therapies that involve the *in vivo* delivery of the Cas9/gRNA system. Much of the effort to treat disease *in vivo* has used Cas9 delivery systems that express the Cas9 protein for prolonged periods of time, such as through a non-integrating AAV vector or even through nanoparticle delivery of mRNA to non-dividing hepatocytes (*8, 10–12*). Use of these viral vectors results in sustained expression of Cas9. In cases where a patient has pre-existing antigen specific T-cells against Cas9, memory T-cells might rapidly expand in response to Cas9 being presented by MHC class I molecules on the cell surface and cytotoxic T-cells will target and clear cells presenting Cas9 on their cell surface(*27*). Furthermore, it is possible to envision that constitutive expression of Cas9 in a large proportion of cells of an organ, for example, the liver, could elicit a cytotoxic T-cell response against Cas9 expressing cells that could result in significant toxicity(*35*). These possible outcomes must be considered seriously, as SaCas9 is the homolog most commonly used for *in vivo* genome editing and here we were able to detect antigen specific T-cells against SaCas9.

Pre-existing adaptive immune responses may be of less concern for *ex vivo* therapies that involve the use of Cas9 to edit cells outside of direct contact with the human immune system. Peptide fragments of any protein delivered into cells *ex vivo* are typically only expressed transiently on cells’ MHC class I molecules, and as such, Cas9 peptides may no longer be present at the cell’s surface at the time of transplantation. While the use of Cas9 *ex vivo*, followed by transplantation of edited cells into a patient, is less of a concern, there remains the currently untested hypothesis that Cas9 may still be present at the cell surface at the time of transplantation and stimulate an immune reaction against edited cells. Therefore, one approach to address the expression of immunogenic Cas9 peptides would be to prolong culture of edited cells and delay their transplantation until the Cas9 peptides were no longer expressed. It may also be possible to either shorten the duration of expression of Cas9 or accelerate the turnover of MHC-bound Cas9. Potential solutions to pre-existing adaptive immunity to Cas9 may include the use of immune suppression or immune depletion to prevent severe cell-mediated responses to Cas9, the use of Cas9 homologs from other bacterial species that do not infect or reside in humans, or the engineering of recombinant Cas9 proteins that escape immune detection. Besides testing of their efficacy in gene editing, exploration of alternatives to SaCas9 or SpCas9 needs to test their immunogenicity in human systems.

In conclusion, our findings raise important new considerations in applying the Cas9/gRNA system to edit human cells for therapeutic purposes. In future work, more sensitive assays can be used to fully understand the pre-existing immune response in humans to Cas9 proteins. We believe our findings will stimulate crucial discussions in the genome editing community about how to most safely and effectively apply the Cas9/gRNA genome editing system for gene therapy in humans.

## Methods

### Cas9 Purification

Recombinant His-tagged SpCas9 protein was expressed from pet28b-Cas9-His in Rosetta 2 cells (EMD Millipore) and Recombinant SaCas9 was cloned from BPK2139 into a pet-28b-His backbone and expressed in Rosetta 2 cells. Bacteria were grown in ZYP-5052 (VWR) at 37C for 12 hours followed by 24 hours at 18C. The bacterial pellet was centrifuged at 6000g for 15 minutes at 4C. The Pellet was re-suspended in lysis buffer (50mM NaH2PO4, 300mM NaCl, 10mM imidazole, 4mM DTT, 5mM Benzamidine, 100mM phenylmethylsulfonyl fluoride, pH 8) and lysozyme was added at 1 mg/ml. After incubation at 4°C for 30 minutes, the lysate was sonicated for six ten second bursts at 200W, with ten second intervals. Lysate was spun at 16,200 g for 1 hr at 4°C and the supernatant was bound to Ni-NTA agarose (Qiagen) at 200rpm, 4°C for 1 hour. The slurry was loaded onto a column and then 50 column volumes of wash buffer with 0.1% tritonX-114 was run over it to remove endotoxin as described previously(*36*) (50mM NaH2PO4, 300mM NaCl, 20mM imidazole, 0.1% TritonX-114, pH 8). The sample was then washed with 20 column volumes of wash buffer (50mM NaH2PO4, 300mM NaCl, 20mM imidazole, pH 8) and eluted with elution buffer (50mM NaH2PO4, 300mM NaCl, 250mM imidazole, pH 8). The eluted SpCas9 and SaCas9 was concentrated using a 100kDa Amicon Ultrafilter (Millipore) and stored in a solution of 10mM Tris-HCl, 150mM NaCl, 50% glycerol.

### Electroporation of Cas9 into CD34+ cells

CD34+ cord blood cells and cord blood serum were provided by the Binns Family Program for Cord Blood research. SpCas9 and SaCas9 was electroporated into one million CD34+ cells in 100ul of T-cell nucleofector solution (Lonza) cells using the Lonza 2b Nucleofector program U-014 (Lonza) at a concentration of 300ug/ml.

### Western Blotting

1 ug of SpCas9 (either His-tag purified or Alt-R SpCas9 (IDT)) and SaCas9 protein as well as cell lysates from 3*10^4 cells were resolved in Laemlli’s sample buffer applied to a 5–15% polyacrylamide gel. Samples were transferred to a polyvinylidene fluoride membrane and blocked with 5% non-fat milk in TBST (50mM Tris-HCl, 150mM NaCl, .05% Tween 20, pH 7.6) for one hour at 4C. Blots were then incubated overnight in TBST with .05% non-fat milk with 1:10 dilution of serum. Blots were then washed five times for five minutes in TBST on a shaker and then incubated with anti-human Fc secondary antibody HRP conjugated at a dilution of 1:5000 for one hour at room temperature. Samples were then washed five times for five minutes in TBST and developed using clarity western ECL substrate (Bio-Rad) and imaged. Cord blood serum was provided by the Binns Family Program for Cord Blood research and adult serum was provided by the Stanford Blood Bank.

### Harvesting, Storage, and Thawing of PBMCs

PBMCs were isolated from apheresis products, from healthy donors obtained from the Stanford Blood Bank, via Ficoll-Paque (density: 1.077 g/mL, GE) density-gradient centrifugation, according to manufacturer’s instructions. Cells were cryopreserved at 50 × 10^6^ cells/mL in CryoStor CS10 (BioLife Solutions) using a rate-controlled container. Cryopreserved PBMCs were thawed for 1 hr at 37°C using thawing medium containing RPMI-1640 (GE, Corning, Live Technologies), 30% human AB serum (Corning), and 10 ug/mL DNAse I (Worthington Biochemical Corporation) and then washed with PBS (Life Technologies, Corning) at 300 × g for 10 min.

### Antigen Stimulation of PBMCs

Cells were incubated at 37°C and 5% CO_2_. For each stimulation, 10^7^ freshly-thawed PBMCs were cultured in 14 mL-plastic culture tubes (Corning) in 1 mL of medium (RPMI-1640 and 10% human AB serum) for 12 hr prior to the addition of antigen. PBMCs were stimulated with 20 ug/mL of either whole purified SpCas9, Alt-R Cas9 (IDT) or SaCas9 protein for 12 hr prior to antigen-specific T-cell isolation. Positive control PBMC samples were stimulated with a viral peptide pool mixture, consisting of protein fragments from human viruses adenovirus 5, BK virus, cytomegalovirus, Epstein-Barr virus, and herpes virus 6, for 4 hr prior to isolation. This peptide pool mixture contained a final concentration of 1 ug/mL of each of: PepTivator AdV5 Penton (Miltenyi Biotec); PepTivator BKV VP2 (Miltenyi Biotec); PepTivator CMV IE-1 (Miltenyi Biotec); MACS GMP PepTivator EBV Select (antigens: proprietary mix of 13 antigens, Miltenyi Biotec), and HHV6 U90 (JPT Peptide Technologies).

### Isolation, Culture, and Restimulation of Antigen-Specific T-cells

Antigen-specific T-cells from each stimulation condition were isolated via the IFN-γ Secretion Assay – Cell Enrichment and Detection Kit (PE), human (Miltenyi Biotec) per manufacturer’s instructions. Prior to magnetic bead labelling, an aliquot of cells was taken for phenotyping (see “Immunophenotyping”). In some experiments, this phenotyping provided an initial IFN-γ secretion profile screen to identify positive donors for a given antigen.

After isolation of antigen-specific T-cells from stimulated PBMCs (Day 0), unselected cells from the column flowthrough were treated with a 25 ug/mL mitomycin-C (Sigma) solution in RPMI-1640 and 10% human AB serum for 30 min and then washed 3 times, according to manufacturer’s instructions (# 130–054–201, Milltenyi Biotec). The mitomyin-C-treated, autologous feeder cells were plated in wells of a 48-well plate (Corning) for 1 hr prior to the addition of antigen-specific T-cells. The antigen-specific T-cells were added, along with 500 uL of growth medium containing X-VIVO 15 (Lonza) and 10% human AB serum. On Day 5, 400 uL of media was replaced with growth medium and 60 U/mL IL-2 (R&D Systems) without disturbing the cells. On Day 8, the Day 5 procedure was repeated and wells were mixed gently. On Days 11 and 14, the Day 5 procedure was repeated. Periodically, an aliquot of cells was taken for phenotyping (see “Immunophenotyping”).

On Day 17, antigen-specific T-cells were washed and then restimulated with fresh, autologous feeder cells in U-bottom 96-well plates (Corning). These autologous feeder cells were prepared from cryopreserved PBMCs, which were thawed, stimulated with antigen, mitomycin-C treated, and washed, as before. Subsequently, intracellular cytokine staining was used to assess surface markers and the production of cytokines (see “Immunophenotyping”).

### Immunophenotyping

Upon initial antigen stimulation, PBMC were stained, according to manufacturers’ instructions, with LIVE/DEAD blue fluorescent marker (Invitrogen) and the following antibodies: IFN-γ PE (Miltenyi Biotec), CD2 FITC, CD3 PerCP/Cy5.5, CD4 BV605, CD8 BV421, CD69 PE-Cy7, CD154 APC-Cy7, CD25 PE-Texas Red, CD62L AF700, CD45RA APC (BioLegend). To conduct a phenotyping assay of the T-cells in culture, the same antibody panel was used, with the replacement of CD19 AF700 and CD14 APC (BioLegend) on appropriate flow cytometry channels.

After restimulation, intracellular cytokine staining (ICS) was used to assess the specificity and potency of the antigen-specific T-cells. Antigen-specific T-cells were also stimulated with DynaBeads Human T-Activator CD3/CD28 (Invitrogen), serving as a positive control for the staining procedure and an indicator for the maximum amount of activation expected from competent cells within a sample. To prepare for ICS, samples were incubated with brefeldin-A (Abcam) for 4 hr prior to the start of the staining procedure. The ICS procedure was conducted per manufacturer’s instructions at 16 hr after restimulation using the Cytofix/Cytoperm Kit (BD). Prior to fixation and permeablization, cells were stained for the surface markers of the antibody panel listed above. Intracellular cytokine production was assessed by incubation with IFN-γ PE (BD), TNF-α AF700, IL-2 APC (BioLegend), accommodating appropriate flow cytometry channels.

## Acknowledgements

MHP gratefully thanks the support of the Amon Carter Foundation and the Laurie Kraus Lacob Faculty Scholar Award in Pediatric Translational Research for this work. DPD thanks the Stanford Child Health Research Institute (CHRI) Grant and Postdoctoral Award for its support of his work. CTC was supported by the CIRM Bridges to Stem Cell Research Program. We thank the Binns Family Program for Cord Blood Research (Stanford University) for providing cord blood sera used in this work. We thank the other members of the Porteus lab for their helpful comments and suggestions.

MHP serves on the Scientific Advisory Board and holds equity in CRISPR Therapeutics but the company had no input into this work.

## Figure Legends

**Supplemental Figure 1:**
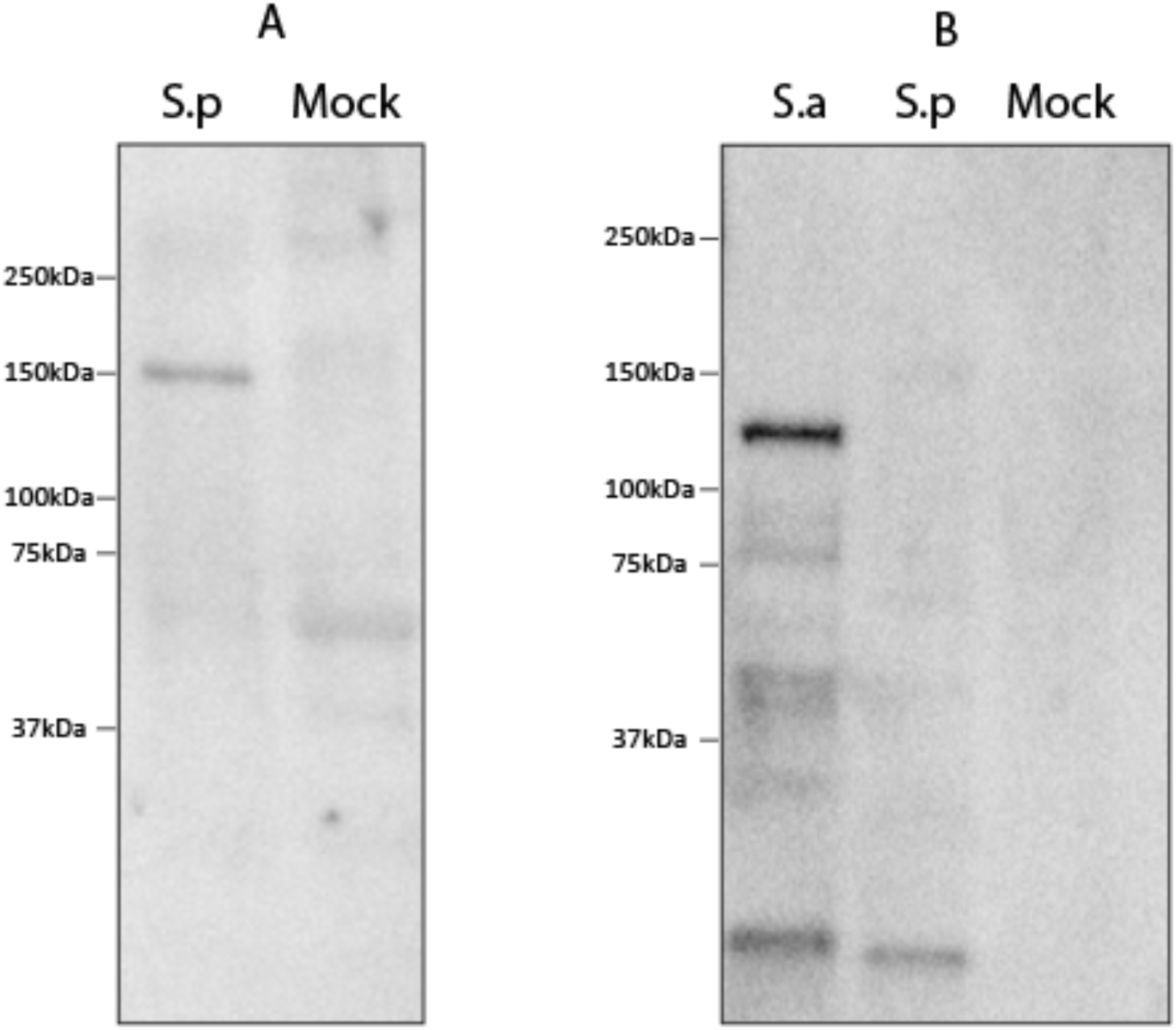
Specificity of Immunoreactivity in Cas9 Expressing CD34+ Cells. CD34+ cells were electroporated with either SaCas9 (S.a) or SpCas9 (S.p) at a concentration of 300ug/ml. Cells were then lysed and the equivalent of 3x10^4^ cord blood derived CD34+ cells were run on a 5–15% gradient gel and blotted. Blots were then probed overnight with Serum from two different human donors (A & B) overnight at a dilution of 1:10 and and anti-human Fc antibody conjugated to HRP was used for detection. Mock denotes CD34+ cell lysates that were not electroporated with Cas9. Immune reactivity was only detected in cells that contained Cas9 protein.

**Supplementary Figure 2:**
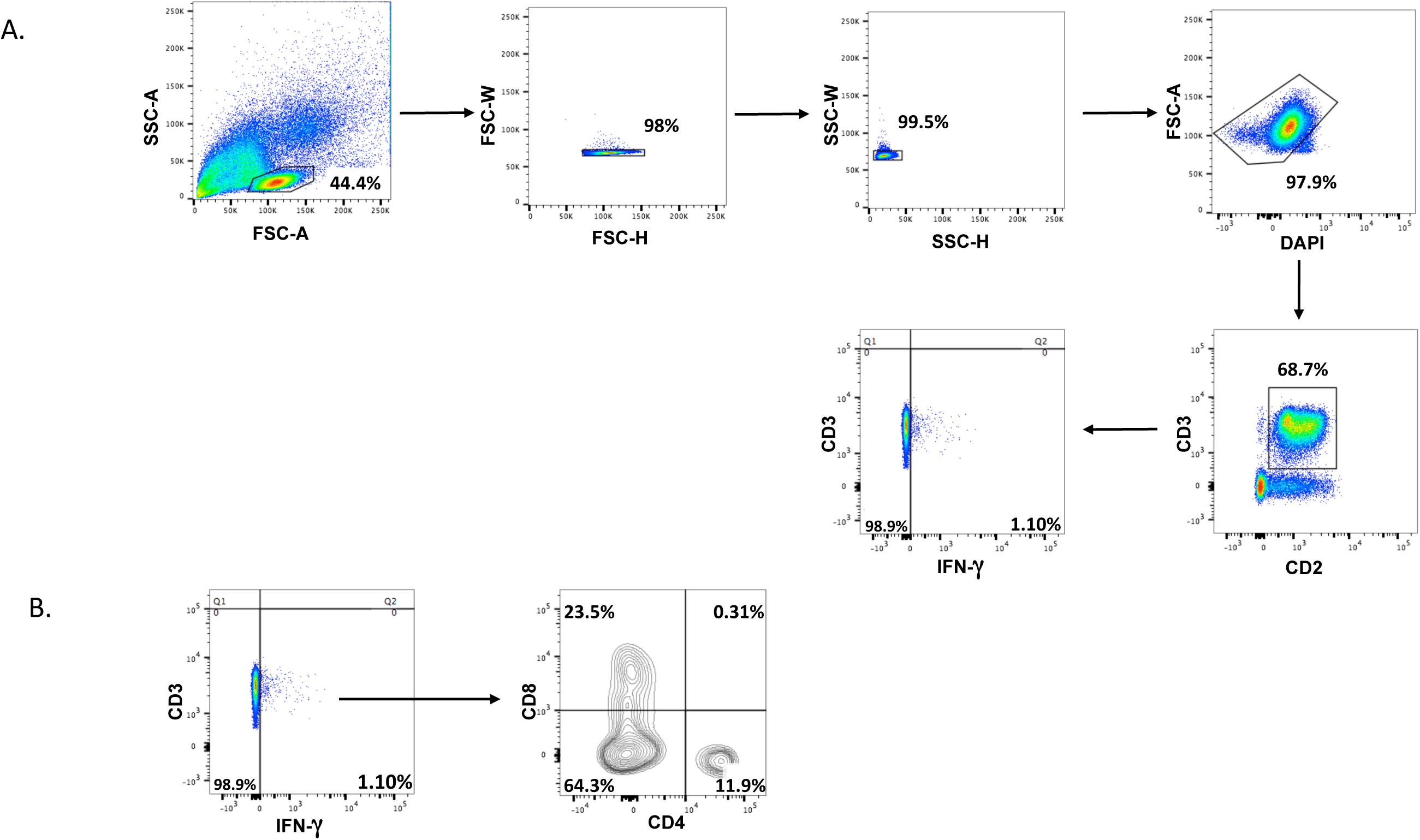
Flow cytometry gating strategy for identifying live, IFN-γ secreting T-cells following cytokine capture system procedure. A) Gating strategy to measure live T-cells producing IFN-γ. B) Gating strategy to determine if these live T-cells producing IFN-γ are CD4+ or CD8+.

**Supplementary Figure 3:**
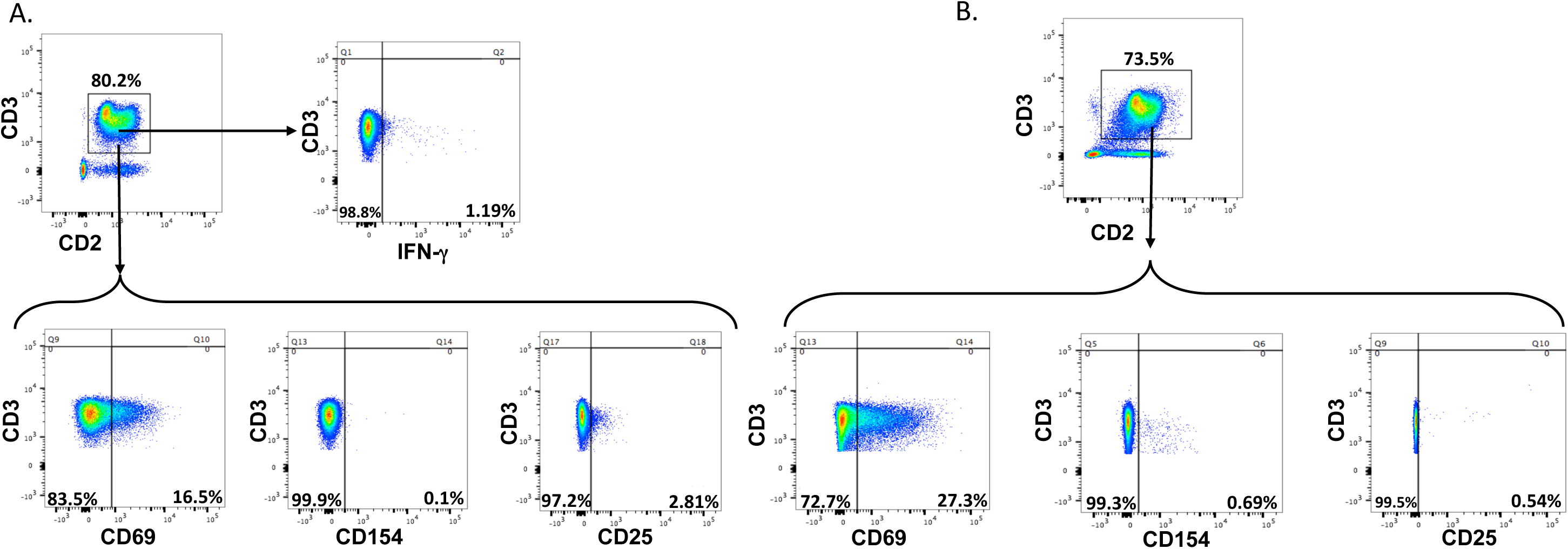
Immunoreactive T-Cells express Markers of Activation. A) Flow cytometry gating of live T-cells after first *in vitro* stimulation, just prior to magnetic isolation step in cytokine capture procedure, to confirm IFN-γ production and to assess activation levels using markers CD69, CD154, and CD25. B) Flow cytometry gating of live, cultured antigen-specific T-cells after re-stimulation with antigen and intracellular cytokine staining (17 days after CCS procedure) to assess of activation levels using markers CD69, CD154, and CD25.

**Supplementary Figure 4:**
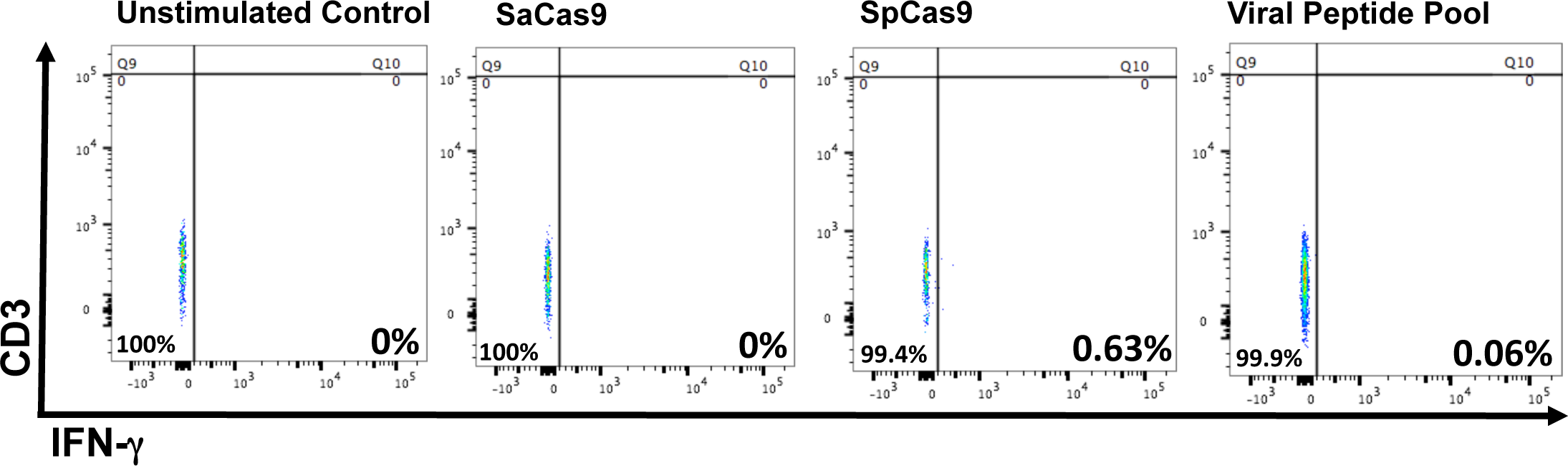
Lack of T-cell Reactivity Derived from Cord Blood. Representative flow cytometry plots identifying negligible levels of T-cells secreting IFN-γ following antigen stimulation and cytokine capture system from CD34-fraction of human cord blood.

**Supplementary Figure 5:**
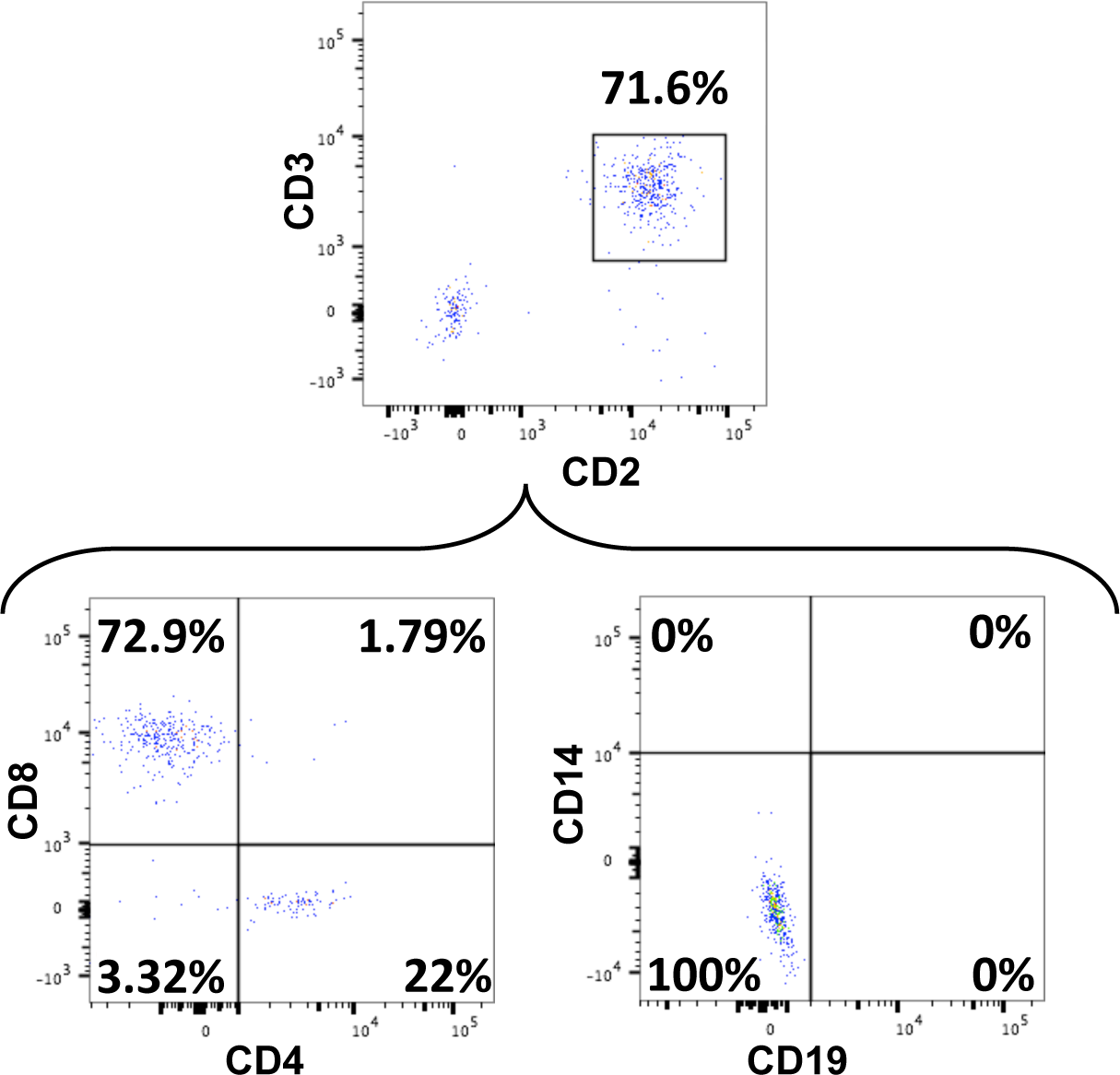
Immunophenotyping of antigen-specific T-cells in culture. Representative immunophenotyping via flow cytometry of live antigen-specific T-cells in culture, in this case, direct against SaCas9, demonstrates the presence CD4+ and CD8+ SaCas9 T-cells and the absence of CD14+ monocytes and CD19+ B-cells.

**Supplementary Figure 6:**
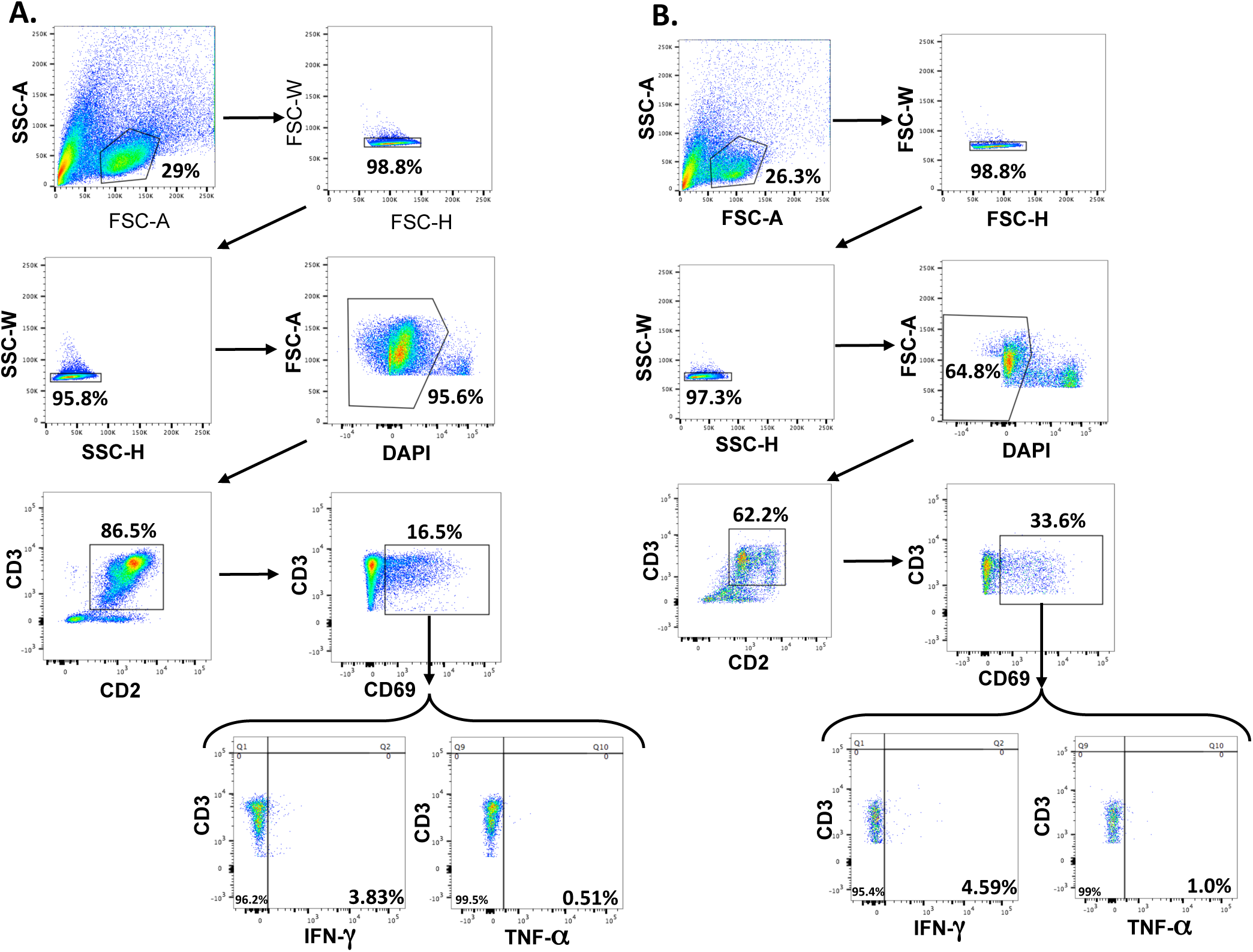
Reactive anti-SaCas9 and anti-viral T-cells express both IFN-γ and TNF-α when re-stimulated with specific antigen. Flow cytometric analyses and gating strategy to determine intracellular IFN-γ and TNF-α production by live, activated CD69+ antigen-specific T-cells upon re-stimulation with specific antigen, following intracellular cytokine staining procedure. Representative for A) anti-SaCas9 T-cells and B) anti-viral T-cells.

**Supplementary Figure 7:**
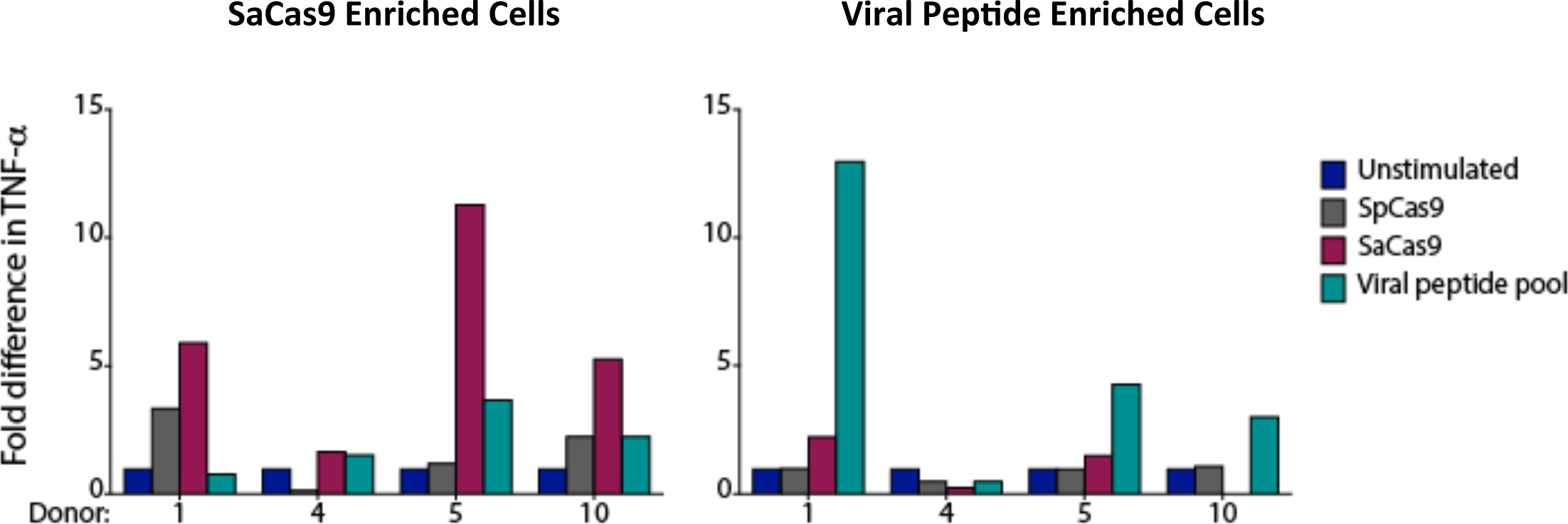
Immunoreactive SaCas9 T-cells Secrete TNF-a upon re-stimulation. Bar graphs showing the fold difference in intracellular TNF-α after cells were re-challenged with antigens after expansion.

**Supplementary Table 1:**
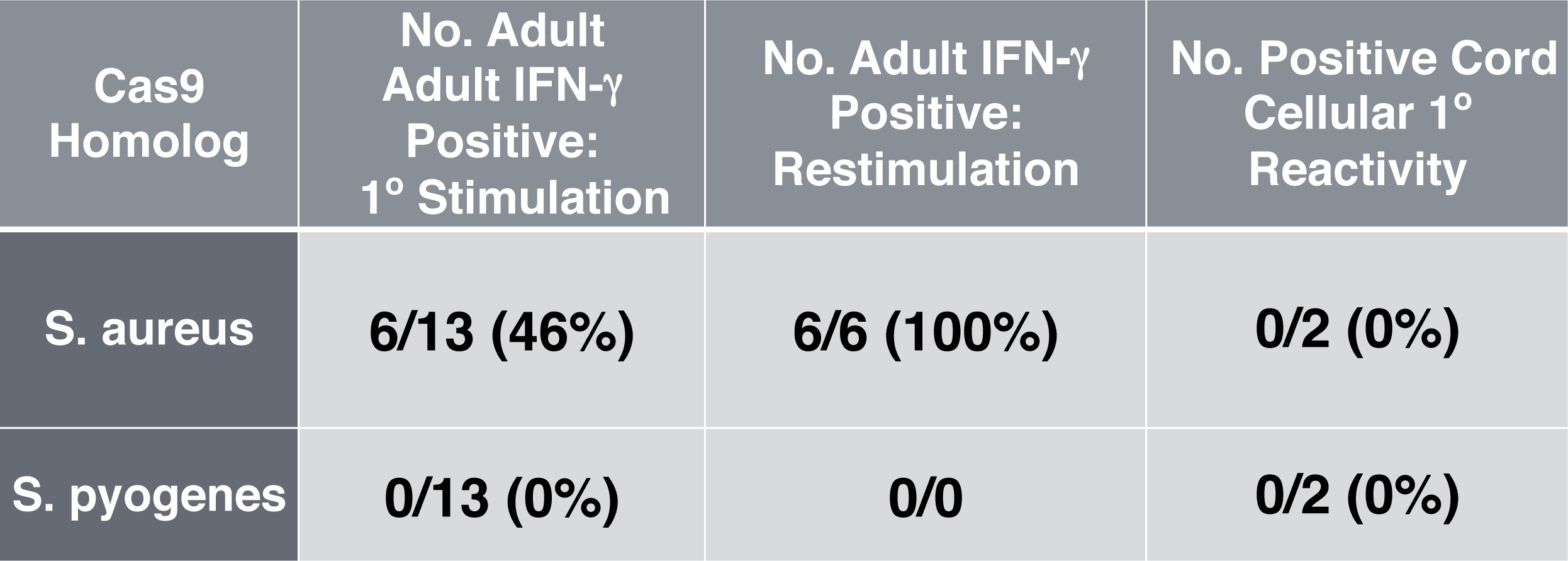
Frequencies at which T-cells were IFN-g Positive upon Stimulation with Cas9 protein

## References

1. L. Cong et al., Multiplex Genome Engineering Using CRISPR/Cas Systems. Science 339, 819 (2013).

2. G. Wang et al., Modeling the mitochondrial cardiomyopathy of Barth syndrome with induced pluripotent stem cell and heart-on-chip technologies. Nat Med 20, 616–623 (2014).

3. M. Jinek et al., A Programmable Dual-RNA-Guided DNA Endonuclease in Adaptive Bacterial Immunity. Science 337, 816 (2012).

4. D. P. Dever et al., CRISPR/Cas9 β-globin gene targeting in human haematopoietic stem cells. Nature 539, 384–389 (2016).

5. J. Eyquem et al., Targeting a CAR to the TRAC locus with CRISPR/Cas9 enhances tumour rejection. Nature 543, 113–117 (2017).

6. M. A. DeWitt et al., Selection-free genome editing of the sickle mutation in human adult hematopoietic stem/progenitor cells. Science Translational Medicine 8, 360ra134 (2016).

7. J. A. Zuris et al., Cationic lipid-mediated delivery of proteins enables efficient protein-based genome editing in vitro and in vivo. Nat Biotech 33, 73–80 (2015).

8. F. A. Ran et al., In vivo genome editing using Staphylococcus aureus Cas9. Nature 520, 186–191 (2015).

9. H. Yin et al., Therapeutic genome editing by combined viral and non-viral delivery of CRISPR system components in vivo. Nat Biotech 34, 328–333 (2016).

10. C. E. Nelson et al., In vivo genome editing improves muscle function in a mouse model of Duchenne muscular dystrophy. Science 351, 403 (2016).

11. M. Tabebordbar et al., In vivo gene editing in dystrophic mouse muscle and muscle stem cells. Science 351, 407 (2016).

12. K. Suzuki et al., In vivo genome editing via CRISPR/Cas9 mediated homology-independent targeted integration. Nature 540, 144–149 (2016).

13. C. Long et al., Prevention of muscular dystrophy in mice by CRISPR/Cas9-mediated editing of germline DNA. Science 345, 1184–1188 (2014).

14. F. D. Lowy, Staphylococcus aureus Infections. New England Journal of Medicine 339, 520–532 (1998).

15. A. L. Roberts et al., Detection of group A Streptococcus in tonsils from pediatric patients reveals high rate of asymptomatic streptococcal carriage. BMC Pediatrics 12, 3 (2012).

16. P. Colque-Navarro, G. Jacobsson, R. Andersson, J.-I. Flock, R. Möllby, Levels of Antibody against 11 Staphylococcus aureus Antigens in a Healthy Population. Clinical and Vaccine Immunology: CVI 17, 1117–1123 (2010).

17. A. Dryla et al., Comparison of Antibody Repertoires against Staphylococcus aureus in Healthy Individuals and in Acutely Infected Patients. Clinical and Diagnostic Laboratory Immunology 12, 387–398 (2005).

18. R. Mortensen et al., Adaptive Immunity against *Streptococcus pyogenes* in Adults Involves Increased IFN-γ and IgG3 Responses Compared with Children. The Journal of Immunology 195, 1657 (2015).

19. J. B. Kolata et al., The Fall of a Dogma? Unexpected High T-Cell Memory Response to Staphylococcus aureus in Humans. The Journal of Infectious Diseases 212, 830–838 (2015).

20. W. L. Chew et al., A multifunctional AAV-CRISPR-Cas9 and its host response. Nat Meth 13, 868–874 (2016).

21. C. S. Manno et al., Successful transduction of liver in hemophilia by AAV-Factor IX and limitations imposed by the host immune response. Nat Med 12, 342–347 (2006).

22. N. J. DePolo et al., VSV-G Pseudotyped Lentiviral Vector Particles Produced in Human Cells Are Inactivated by Human Serum. Molecular Therapy 2, 218–222 (2000).

23. F. Mingozzi et al., CD8+ T-cell responses to adeno-associated virus capsid in humans. Nat Med 13, 419–422 (2007).

24. J. R. Mendell et al., Dystrophin Immunity in Duchenne’s Muscular Dystrophy. New England Journal of Medicine 363, 1429–1437 (2010).

25. J. M. Wilson, Lessons learned from the gene therapy trial for ornithine transcarbamylase deficiency. Mol Genet Metab 96, 151–157 (2009).

26. F. Mingozzi, K. A. High, Immune responses to AAV vectors: overcoming barriers to successful gene therapy. Blood 122, 23–36 (2013).

27. E. W. Hewitt, The MHC class I antigen presentation pathway: strategies for viral immune evasion. Immunology 110, 163–169 (2003).

28. K. M. Smith et al., Th1 and Th2 CD4^+^ T Cells Provide Help for B Cell Clonal Expansion and Antibody Synthesis in a Similar Manner In Vivo. The Journal of Immunology 165, 3136 (2000).

29. J. D. M. Campbell et al., Rapid detection, enrichment and propagation of specific T cell subsets based on cytokine secretion. Clinical & Experimental Immunology 163, 1–10 (2011).

30. J. F. Möller, B. Möller, B. Wiedenmann, T. Berg, E. Schott, CD154, a marker of antigen-specific stimulation of CD4 T cells, is associated with response to treatment in patients with chronic HCV infection. Journal of viral hepatitis 18, e341–e349 (2011).

31. P. E. Simms, T. M. Ellis, Utility of flow cytometric detection of CD69 expression as a rapid method for determining poly- and oligoclonal lymphocyte activation. Clinical and Diagnostic Laboratory Immunology 3, 301–304 (1996).

32. B. Fazekas De St. Groth, A. L. Smith, C. A. Higgins, T cell activation: in vivo veritas. Immunol Cell Biol 82, 260–268 (2004).

33. S. E. Christmas, A. Meager, Production of interferon-gamma and tumour necrosis factor-alpha by human T-cell clones expressing different forms of the gamma delta receptor. Immunology 71, 486–492 (1990).

34. K. Kuwano, T. Kawashima, S. Arai, Antiviral effect of TNF-α and IFN-γ secreted from a CD8+ influenza virus-specific CTL clone. Viral immunology 6, 1–11 (1993).

35. Y. Yang, J. M. Wilson, Clearance of adenovirus-infected hepatocytes by MHC class I-restricted CD4+ CTLs in vivo. The Journal of Immunology 155, 2564 (1995).

36. P. Reichelt, C. Schwarz, M. Donzeau, Single step protocol to purify recombinant proteins with low endotoxin contents. Protein Expression and Purification 46, 483–488 (2006).

